# Private QTLs and the Genetic Architecture of Hierarchical Size Traits: From Body Size to Sex-Specific Plasticity

**DOI:** 10.1101/2025.10.30.685544

**Authors:** Isabelle Vea, Carlos Molina, S. Wilcox Austin, W. Anthony Frankino, Alexander W. Shingleton

## Abstract

Sex-specific size plasticity (SSP) is the phenomenon whereby the size of one sex is more environmentally sensitive than the other, and is thought to underlie the developmental regulation and evolution of sexual size dimorphism (SSD). Sex-specific plasticity is a higher order phenotype that emerges due to the effect of the environment and sex on core growth regulatory mechanisms. Genetic variation in SSP necessarily requires sex- and environment-specific variation in growth, yet the developmental-genetic mechanisms enabling such context-dependent size variation remain poorly understood. Using a genome-wide association study (GWAS) and functional validation in *Drosophila melanogaster*, we dissected the genetic architecture of body size, size plasticity, SSD, and SSP across 196 isogenic lineages. We find that each phenotype is governed by largely non-overlapping sets of loci, with most candidate variants lying outside canonical growth pathways. Instead, size trait are shaped by “private QTLs”, whose effects are limited to specific sex, trait, or environmental contexts. Functional knockdown of selected candidate genes for SSP revealed that while most did not affect SSP directly, many influenced body size, SSD, and size plasticity, in a manner consistent with their nested phenotypic relationships. Together, our results suggest that context-dependent alleles in genes peripheral to core growth regulatory pathways drive variation in SSD and SSP, offering a mechanistic explanation for their evolutionary lability and highlighting the role of private QTLs in structuring complex trait architecture.

**Significance Statement:** Sexual size dimorphism (SSD), the differences in body size between females and males, is highly variable among species and yet the developmental genetic mechanisms that generate it and drive its evolution remain elusive. Our work reveals a modular genetic architecture underlying SSD in *Drosophila melanogaster*, showing that variation in body size, its sex-specific plasticity, and the resulting SSD are controlled by largely non-overlapping, context-dependent loci. This finding provides critical insight into how complex morphological differences between the sexes are genetically organized, allowing for the flexible and independent evolution of hierarchical traits.

## Introduction

Apart from the possession of different reproductive organs, Sexual Size Dimorphism (SSD) is perhaps the most common phenotypic difference between females and males in animals. In most arthropods and poikilothermic vertebrates, females are the larger sex (1–3), whereas the reverse pattern is typical in mammals and birds (4, 5), although see (6). SSD is extremely evolutionarily labile, changing dramatically among even closely related species (7–10). Over a century of research has focused on the selective pressures that generate SSD and that account for differences in SSD among species (11–13).

Nevertheless, we have a comparably poor understanding of the developmental genetic mechanisms these selective pressures target to generate differences in body size between females and males, representing a considerable gap in our knowledge of the evolution of SSD. This gap is widened because SSD cannot (usually) be measured in an individual, making it challenging to describe the genetic variation in SSD upon which selection can act.

Our previous work circumvented the challenge of characterizing genetic variation in SSD among individuals by measuring SSD in 196 isogenic *Drosophila* lineages, each with a unique genotype (14). We demonstrated substantial genetic variation in sexual size dimorphism (SSD), primarily driven by greater genetic variation in body size among females compared to males. Although we observed a general trend of increasing SSD with sex-averaged body size across lineages (contrary to Rensch’s Rule (15)), the majority of SSD variation was independent of sex-averaged body size. Instead, SSD exhibited a strong genetic correlation with nutritionally induced sex-specific size plasticity (here referred to as SSP), the phenomenon where body size is more sensitive to variation in developmental nutrition in one sex. It is a truism that nutritional SSP generates SSD, at least under some dietary conditions. Our data suggest that females from lineages with heightened SSD grew substantially larger than males when both sexes were well-fed relative to when they were malnourished (14).

These findings support the *condition dependence hypothesis* of sexual dimorphism (9, 16–20), which predicts that when one sex experiences stronger directional selection and greater marginal fitness gains from increased trait size, it will invest more in trait growth under better conditions. In insects like *Drosophila*, fecundity selection drives stronger selection on female body size, as reproductive output scales positively with size (21, 22). As a result, females enjoy greater marginal fitness gains more from increased size than males, and thus allocate more to growth under high nutrition, producing female-biased SSP for body size (14).

The link between SSP and SSD is further supported by developmental evidence showing that, in *Drosophila*, SSD appears to be regulated by the insulin/insulin-lie growth factor signaling (IIS) pathway (23, 24). The IIS is the major regulator of growth with respect to nutrition in all animals (25, 26). SSD is dependent on functional IIS and is lost or reduced in malnourished flies or those with mutations in the IIS pathway (24, 27). Further, well-fed females appear to have higher IIS activity than well-fed males (22), consistent with their larger size. The result is that female body size is more insulin-sensitive than male body size, functionally accounting for the observed SSP.

A compelling hypothesis, therefore, is that variation in SSP and SSD is coregulated by genetic variation in the signaling pathways that regulate growth in response to nutrition, in particular the IIS pathway. This hypothesis, however, presents a puzzle. SSP is a *higher-order phenotype*; that is, it emerges from the effect of diet on SSD, or, conversely, from the effect of sex on size plasticity. Further, SSD and size plasticity are themselves higher-order phenotypes that emerge due to sex-specific and diet-specific effects on growth. Thus, genetic variants that influence SSP, SSD, and size plasticity necessarily have highly contextual effects, impacting body size only in specific nutritional environments and/or in one sex. It is difficult to reconcile this pattern of trait-specific genetic effects with variation in core growth-regulatory genes, such as those in the IIS pathway, which are employed broadly across tissues, environments, and sexes. Changes in these central pathways are expected to produce widespread pleiotropic consequences affecting growth in multiple tissues, in both sexes, and under different nutritional conditions. This raises a central, paradoxical question: How can the genetic architecture underlying sex- and environment-specific phenotypes be built from components that, by their very nature, act globally on growth?

To answer this question, we conducted a genome wide association study (GWAS) and subsequent functional validation experiments to characterize the genetic architecture that underlies variation in body size, size plasticity, SSD, and SSP using 196 isogenic *Drosophila* lineages. Surprisingly, we find almost no overlap in the alleles that underlie variation for each phenotype, with almost all candidate alleles lying in genes outside canonical growth-regulatory and nutrient-sensing pathways. Rather, standing genetic variation in these traits is driven primarily by private QTLs: loci in genes that peripherally regulate general growth, but whose allelic effects are limited to specific trait, sex, or environment combinations. These private QTLs explain how hierarchical traits can evolve independently, and why body size, size plasticity, SSD, and SSP can be labile even when regulated by a common developmental mechanism.

## Results

### Genetic-Basis of Variation in Body Size, Sexual Size Dimorphism and Sex Specific Plasticity

We conducted a GWAS on multiple body size traits, including fed and starved female and male body size 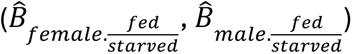, fed and starved sex-averaged body size 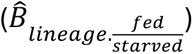, fed and starved sexual size dimorphism (male and female body size plasticity (Δ_*female*_, Δ_*male*_), sex-averaged plasticity ( Δ_*lineage*_), and sex-specific plasticity (SSP). Each analysis followed a two-step approach. First, we performed a rapid GWAS using the DGRP2 pipeline on Best Linear Unbiased Predictions (BLUPs) per lineage to pre-screen SNPs. Second, we applied a more computationally intensive mixed linear model (MLM) GWA using all individual phenotypic data, restricted to the 10,000 SNPs with the lowest *p*-values from the DGRP2 GWAS. The MLM GWAS is both more powerful and less prone to Type I error than the DGRP2 GWAS, as it incorporates phenotypic variation both within and between isogenic lineages. Using the results of our MLM GWAS we then conducted a Versatile Gene-based Association Study (VEGAS) (28) to test whether multiple SNPs within the same gene carried an aggregate signal of association. We refer to SNPs with *p*< 1×10⁻⁵ in our MLM GWA as *candidate SNPs*, and genes with *p*< 1×10⁻⁵ in our VEGAS as *candidate genes*. A list of GWAS and VEGAS hits (*p*< 1×10⁻⁵) is provided in **Supplementary Table S1**, and the complete results of the DGRP2 GWAS, MLM GWAS and VEGAS is provided in **Supplementary Table S2** (available at DOI: 10.5061/dryad.8sf7m0d36).

Consistent with previous GWAS elucidating the genetic architecture of body size traits in *Drosophila* (29, 30) our MLM GWAS found a moderate (10–40) number of SNPs associated with variation in body size, SSD, and SSP, corresponding to a similar number of candidate genes from our subsequent VEGAS (**Figure 1**). In general, genotypes with the major allele for the candidate SNPs for SSP showed a reduction in SSD upon starvation, which is the general pattern in *Drosophila* (14), while lineages with the minor allele tended to either reduce or reverse this pattern (**Figure 1C**).

**Figure 1.**
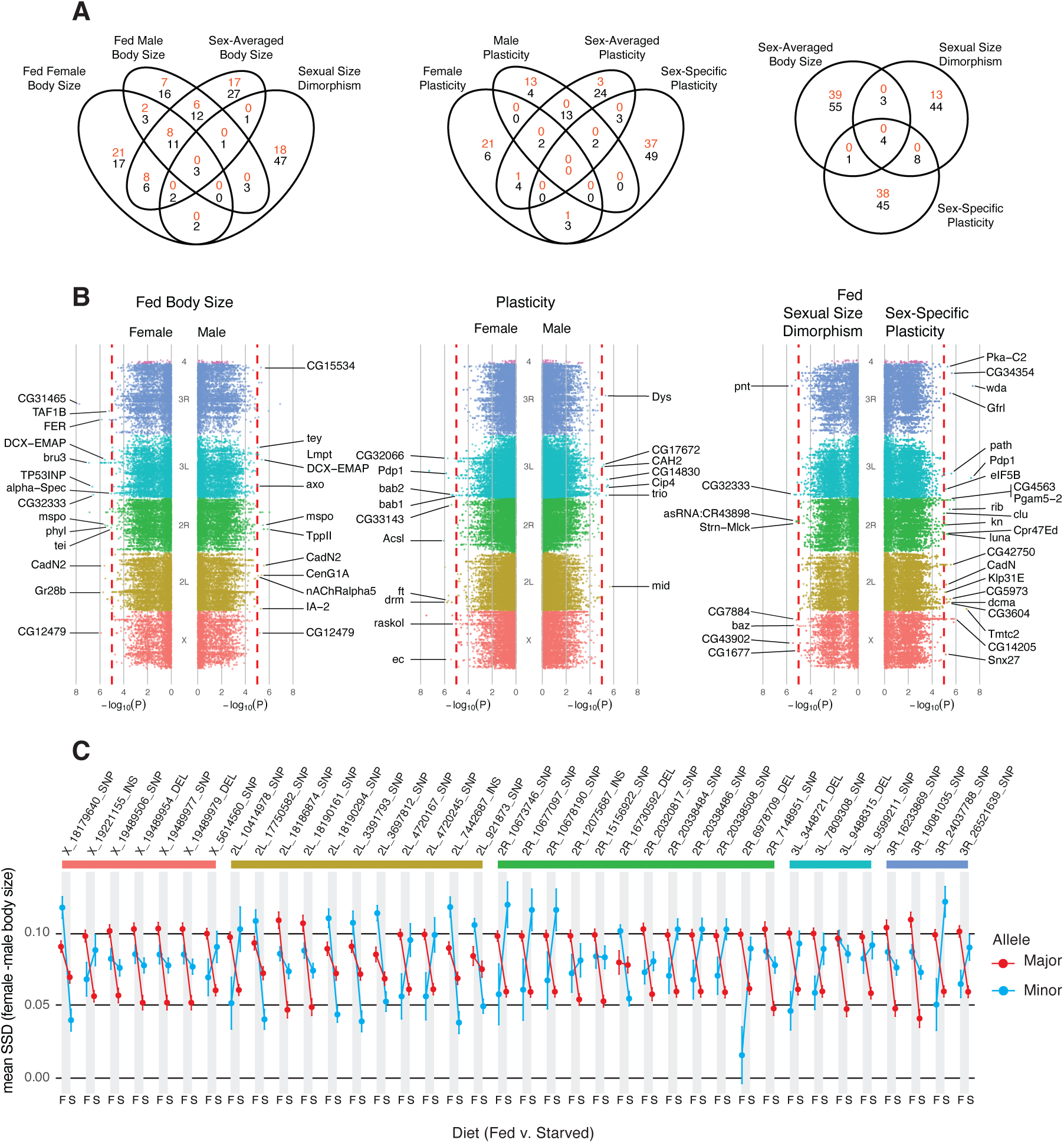
Genetic variants affecting body size, SSD, plasticity and sex-specific plasticity. (A) Overlap of candidate SNPs (red) and genes (black) for different size-related phenotype. Candidate SNPs were identified by MLM GWAS and candidate genes were identified by VEGAS. (B) Manhattan plots of SNPs tested using MLM GWAS, with candidate SNPs labelled according to the gene they lie in/adjacent to (Supplementary Tables S1 and S2).(C) Mean and 95% confidence intervals for SSD in fed (F) and starved (S) flies for the major (red) and minor (blue) allele of candidate SNPs for SSP. A change in SSD with diet indicate sex-specific plasticity. SNPs are arranged according to their genomic position.

In an earlier study we found a weak but significant genetic correlation between female body size and SSD among DGRP lineages, accounting for 17% of the variation in SSD, but not between male body size and SSD. Correspondingly, we detected a weak but significant positive correlation between sex-averaged body size and SSD, contributing to 10% of the variation in SSD. We found almost no overlap, however, between either candidate SNPs or genes for SSD and either male, female or sex-averaged body size in either fed flies (**Figure 1A, B**) or starved flies (**Supplementary Figure S1A**). Further, there was almost no overlap between candidate SNPs and genes across environmental conditions (fed v. starved, **Supplementary Figure S1B**).

A GO analysis of candidate genes for female, male and sex-averaged body size under both fed and starved conditions revealed enrichment for processes tied to neuronal and epithelial morphogenesis, as well as imaginal disc morphogenesis (**Supplementary Table S3**). Enriched cellular components centered on junctional and synaptic regions of the plasma membrane, and molecular functions emphasize signaling and adhesion (e.g., growth-factor receptor binding and neurotransmitter receptor activity). A KEGG analysis suggested possible enrichment for purine metabolism and neuroactive ligand–receptor signaling (at *q*<0.1), although the signal was weak (**Supplementary Table S3**)

We had also previously observed a genetic correlation between SSD and SSP among the lineages, with SSD explaining 32% of the genetic variation in SSP. However, we again found almost no overlap between the candidate SNPs and genes associated with variation in SSD and SSP (**Figure 1A, B**). Similarly, we found almost no overlap between the candidate SNPs or genes for SSP and either male, female or sex-averaged plasticity (**Figure 1A, B**). A GO analysis of all the candidate genes for body size plasticity (male, female, sex-averaged and sex-specific) found enrichment for neurodevelopmental terms, such as neuron and axon projection development, axon guidance/ogenesis, and dendrite morphogenesis, as well as cell-adhesion and cytoskeletal processes (cell–cell adhesion, junction organization, actin-based pathways) (**Supplementary Table S3**). Several imaginal disc morphogenesis and development terms also appeared, with particularly strong enrichment for the semaphorin-plexin signaling pathway, suggesting regulators of body size plasticty are tightly linked to neuronal wiring and developmental morphogenesis. A KEGG analysis of these genes highlighted potential enrichment for developmental signaling pathways, particularly Wnt, Hippo, Hedgehog, and dorso-ventral axis formation (at *q*<0.1), though most categories were represented by only two to four genes. A KEGG analysis on SSP candidate genes alone provided slightly stronger support for the involvement of core developmental and growth-regulatory pathways, especially Wnt, Hedgehog, Hippo, and circadian rhythm (**Supplementary Table S3**). Other pathways, including metabolism and nutrient-signaling (mTOR and FOXO), were also detected but with weaker support and higher *q*-values

Our study was conducted explicitly to elucidate the genetic architecture of the nutritional plasticity of body size in females and males. We were therefore surprised by the paucity of known growth regulators among the candidate genes in general, and nutritional-regulators of growth specifically. One plausible explanation is that core growth-control genes are constrained (often essential), harboring predominantly small-effect SNPs that do not yield sufficiently low *p*-values to meet our criteria for candidate SNPs. To test whether predefined gene sets nonetheless show aggregate signal, we conducted a gene-set permutation analysis: For each gene in the set (330 growth regulatory genes and 70 nutrient-signaling genes, **Supplementary Table S4**), we identified the SNP with the strongest association (lowest *p*-value) and then averaged these minimum values across the gene set. We then compared this index to a null distribution obtained by repeatedly sampling equal-sized, functionally unrelated gene sets from the genome. We applied this analysis to the results of the MLM GWAS of SSP, SSD, sex-averaged plasticity and sex-averaged body size. We found that growth-regulatory genes showed significantly stronger aggregate association than expected by chance for SSP, SSD, sex-averaged plasticity, whereas nutrient-signaling genes did not show similar enrichment, and sex-averaged body size was not enriched for either of the gene sets (**Table 2**).

**Table 1:** Results of permutation test for aggregate GWAS signal among SNPs for growth-regulatory and nutrient-signaling genes.

| Phenotype | Pathway<sup>1</sup> | Number of genes<sup>2</sup> | Observed mean $p$-value<sup>3</sup> | Null mean $p$-value<sup>4</sup> | Null SD<sup>5</sup> | Percentile<sup>6</sup> |
| --- | --- | --- | --- | --- | --- | --- |
| Sex-averaged body size<br>($\hat{B}_{lineage..fed}$) | Growth | 161 | 0.2093 | 0.2440 | 0.0234 | 6.63 |
|  | Nutrient | 24 | 0.2560 | 0.2432 | 0.0619 | 59.50 |
| Sex-averaged plasticity<br>($\Delta_{lineage}$) | Growth | 180 | 0.1935 | 0.2374 | 0.0218 | **2.02** |
|  | Nutrient | 29 | 0.2556 | 0.2377 | 0.0553 | 63.99 |
| SSD<br>($SSD_{fed}$) | Growth | 160 | 0.1666 | 0.2324 | 0.0232 | **0.18** |
|  | Nutrient | 24 | 0.2012 | 0.2324 | 0.0612 | 31.98 |
| SSP | Growth | 164 | 0.1625 | 0.2252 | 0.0232 | **0.26** |
|  | Nutrient | 28 | 0.2606 | 0.2254 | 0.0557 | 74.09 |

<sup>1</sup> Target pathways. Growth-signaling pathways comprise 330 genes, nutrient-signaling pathways comprise 70 genes.
<sup>2</sup> Number of genes containing SNPs tested by MLM GWAS.
<sup>3</sup> Mean $p$ -value for SNPs in growth-regulatory and nutrient-signaling genes.
<sup>4</sup> Mean $p$ -value for SNPs across a randomized set of genes of the same size.
<sup>5</sup> The standard deviation of null mean $p$ -value
<sup>6</sup> The percent of means with a value lower than the observed mean $p$ -value in the null distribution. Significant percentages (<5%) are shown in **bold**.

**Table 2:** Details of candidate genes subject to functional validation.

| Gene ID | FBgn | Function | MLM GWAS<sup>1</sup> | DGRP2 GWAS<sup>2</sup> | Validation<sup>53</sup> |
| --- | --- | --- | --- | --- | --- |
| **Chromatin & transcription regulation** |  |  |  |  |  |
| *Wda* | FBgn0039067 | Histone acetylation complex subunit | **3.92x10<sup>-8</sup>** | **2.93x10<sup>-7</sup>** | **S\*D\*T** |
| *Pdp1* | FBgn0016694 | Circadian and muscle transcription factor | **5.35x10<sup>-8</sup>** | **3.74x10<sup>-8</sup>** | **S\*D\*T** |
| *Luna* | FBgn0040765 | Chromatid separation transcription factor | **4.47x10<sup>-6</sup>** | **5.62x10<sup>-6</sup>** | NS |
| *Kn* | FBgn0001319 | Muscle and wing transcription factor | **6.28x10<sup>-6</sup>** | 1.43x10<sup>-4</sup> | **D\*T & S\*T** |
| **RNA & gene silencing** |  |  |  |  |  |
| *CG34354* | FBgn0085383 | Nuclear mRNA splicing regulator | **2.20x10<sup>-6</sup>** | **1.33x10<sup>-6</sup>** | NS |
| *qin* | FBgn0263974 | piRNA pathway silencing | 1.30x10<sup>-4</sup> | 1.33x10<sup>-4</sup> | NS |
| **Signal transduction** |  |  |  |  |  |
| *Pka-C2* | FBgn0000274 | cAMP-dependent protein kinase | **5.02x10<sup>-6</sup>** | 4.11x10<sup>-5</sup> | T |
| *Wnt10* | FBgn0031903 | Wnt signaling protein | 2.67x10<sup>-3</sup> | **7.25x10<sup>-6</sup>** | **D\*T** |
| **Enzymes & metabolism** |  |  |  |  |  |
| *CG14205* | FBgn0031034 | Non-amino-acyl transfer enzyme | **1.41x10<sup>-6</sup>** | 1.36x10<sup>-5</sup> | NS |
| *Pgam5-2* | FBgn0035004 | Mitochondrial fission phosphatase | **4.70x10<sup>-6</sup>** | **5.11x10<sup>-6</sup>** | **D\*T** |
| *eIF5B* | FBgn0026259 | Translation initiation GTPase | **6.86x10<sup>-6</sup>** | **3.06x10<sup>-6</sup>** | NS |
| *CG32333* | FBgn0052333 | Lipid metabolism enzyme | 2.59x10<sup>-5</sup> | 1.64x10<sup>-5</sup> | **D\*T** |
| *CG12065* | FBgn0030052 | Nucleoside metabolic enzyme | 5.98x10<sup>-5</sup> | **4.57x10<sup>-6</sup>** | **D\*T** |
| *Hn* | FBgn0001208 | Serotonin and tyrosine hydroxylase | 3.89x10<sup>-4</sup> | **8.76x10<sup>-6</sup>** | NS |
| *CARPB* | FBgn0052698 | Hydro-lyase enzyme | 2.38x10<sup>-2</sup> | **5.15x10<sup>-6</sup>** | NS |
| *CG3604* | FBgn0031562 | Extracellular serine protease inhibitor | **2.20x10<sup>-6</sup>** | **1.33x10<sup>-6</sup>** | **D\*T** |
| **Cell adhesion & structural proteins** |  |  |  |  |  |
| *CadN* | FBgn0015609 | Calcium-dependent cell adhesion | **8.36x10<sup>-6</sup>** | 3.06x10<sup>-5</sup> | **S\*T** |
| *BeatVII* | FBgn0250908 | Heterophilic cell adhesion | 2.17x10<sup>-5</sup> | **5.50x10<sup>-6</sup>** | **D\*T & S\*T** |

<sup>1</sup> Lowest *p*-value among gene-specific SNPs in MLM GWAS.
<sup>2</sup> Lowest *p*-value among gene-specific SNPs in DGRP2 GWAS.
<sup>3</sup> Significant (*p*<0.05) effects from functional analysis of gene knockdown (KD) on size. S = sex (male v. female), D = diet (fed v. starved), T = type (experimental v. control). S\*D\*T = KD affects SSP, S\*T = KD affects SSD, D\*T = KD affects plasticity, T = KD affects body size. In all cases, there was a significant effect of sex and diet on body size independent of type.

We also conducted a VEGAS on genes from the same gene sets, to determine whether SNPs within growth-regulatory and nutrient-signaling genes may be associated in aggregate with variation in SSP, SSD and sex-averaged body size. No nutrient-signaling gene had a *p*-value<1×10⁻⁵ in our VEGAS analysis, and only 16 out of 330 growth-regulatory genes were below this threshold for SSP, SSD and sex-averaged body size collectively (**Supplementary Tables S5**).

### Validity of a Two-Step GWAS

Limiting the MLM GWAS to only the top 10,000 SNPs might exclude additional true positives; that is, SNPs with relatively high *p*-values in the DGRP2 GWA but low *p*-values in the MLM GWA. To assess this, we ranked DGRP2 SNPs by their *p*-values and plotted the cumulative number of SNPs with MLM *p*-values < 1×10⁻⁵ as a function of rank. For most traits, this accumulation plateaued by ∼3,000 SNPs, suggesting that the top 10,000 DGRP2 SNPs provided a sufficiently inclusive set for downstream MLM analysis (**Supplementary Figure S2**). The exception was sex-averaged plasticity, where the accumulation plateaued at ∼9,000 SNPs indicating we may have missed a small number of low *p*-values in our MLM GWAS. As a second test of our two-step approach, we applied the MLM GWAS to random subset of 10,000 SNPs not in the top 10,000 to assess if any true positives were missed. In no case did we observe a SNP with a *p*-value < 1×10⁻⁵ in our MLM GWAS on this subset (**Supplementary Table S1**)

### Functional Validation of QTLs for SSP

To functionally validate our GWAS, we focused on candidate genes that were associated with variation in SSP. We further refined our validation to genes with available transgenic RNAi constructs that allow us to knockdown gene expression. In each case we used *act-GAL4* to knockdown gene expression ubiquitously, and compared the effect on sex-specific plasticity relative to control flies reared under the same conditions. We tested eleven candidate genes for SSP regulation based on our MLM GWAS and five based on our DGRP2 GWAS. We also tested two genes that did not pass the threshold for either the MLM or DGPR2 GWAS, one of which was just above the threshold *p-value* and one of which was an order of magnitude above the threshold *p-value* (**Table 3**).

**Table 3.**
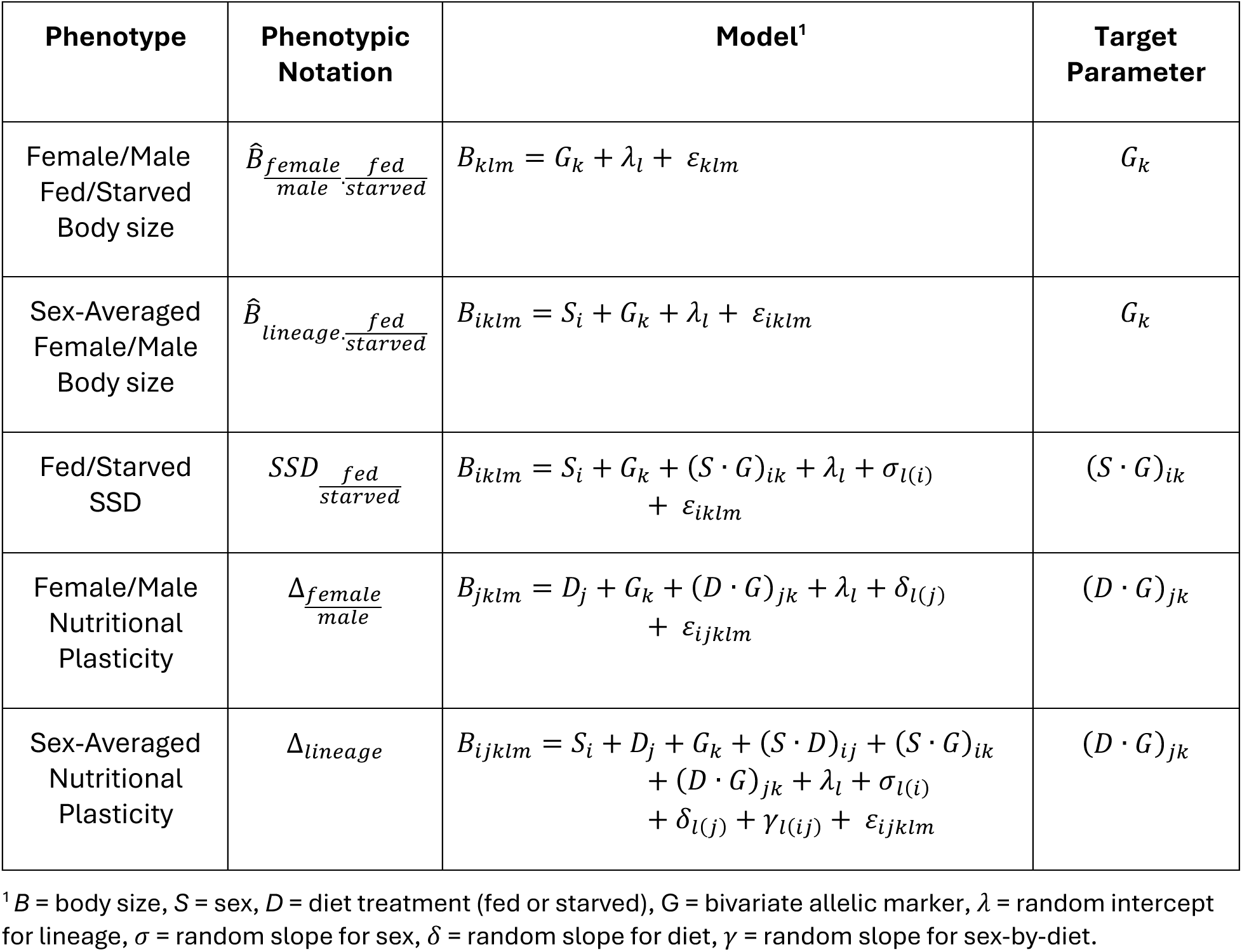
Mixed linear models used to conduct GWA analysis for each phenotype.

Of our candidate genes, only two showed significant effects on SSP when ubiquitously knocked down (**Table 3**, **Figure 2**): *wda*, which encodes a component of the SAGA chromatin-modifying complex essential for histone acetylation and transcriptional regulation, and *Pdp1*, which encodes a PAR-domain bZIP transcription factor that regulates gene expression in circadian clock neurons and muscle development.

**Figure 2.**
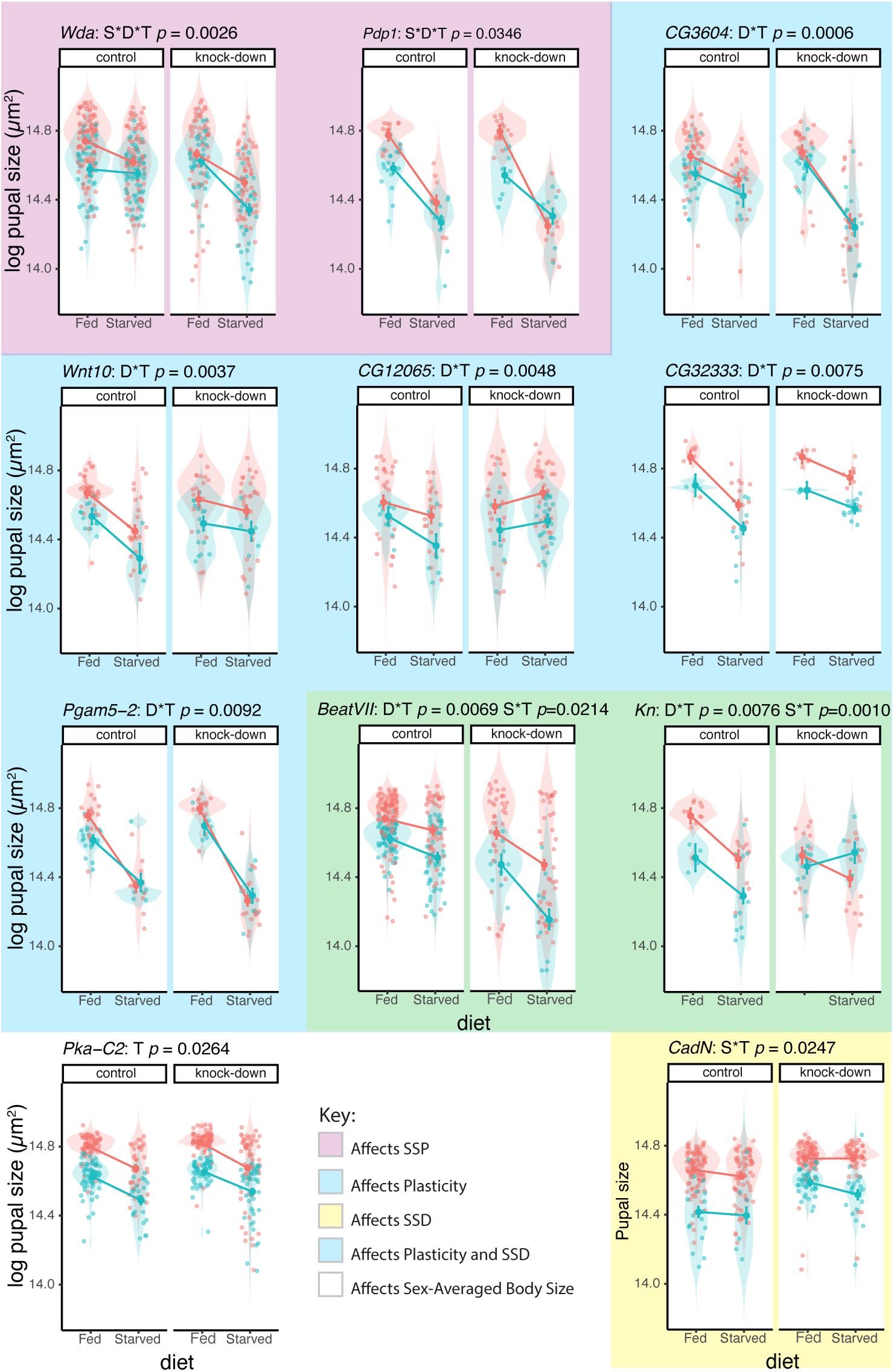
Functional validation of candidate genes for sex-specific plasticity (SSP). Each panel shows results for an individual gene. Panel titles give the gene name and the *p*-values for the effects of gene knockdown on SSP (S·D·T), plasticity (D·T), sexual size dimorphism (S·T), and overall body size (T) (see Table 3). If a higher-order interaction is significant, lower-order tests are not shown. Violin plots show the distribution of pupal size for each treatment group (control vs. knockdown; fed vs. starved). Large points represent fitted means from the model, and error bars are 95 % confidence intervals.

A change in SSP, however, necessarily involves a change in SSD (at least in one environmental condition), a change in plasticity (at least in one sex) and a change in body size (at least in one environmental condition and sex). We found that knockdown of many of the candidate genes did not significantly affect SSP but nevertheless affected SSD, plasticity and/or body size. Overall, 68% of the candidate genes were validated for their effects on body size, plasticity, SSD and/or SSP. Of the remaining two genes that were just above the MLM or DGRP2 *p*-value threshold for SSP, knockdown of one (*CG32333*) was found to affect sex-averaged nutritional plasticity, while knockdown of the other (*eIF5B*) had no detectable affect.

## Discussion

### Developmental Regulation of SSD and SSP

Sexual size dimorphism (SSD) is evolutionarily widespread but highly labile in its expression, varying both in its direction and its magnitude among closely related species and populations. Over a century of research has generated numerous theories as to the ultimate selective pressures that generate SSD. However, we have very little knowledge of the proximate developmental mechanisms that are the target for these selective pressures. Establishing which mechanisms underlie expression and variation in SSD is necessary if we are to completely evaluate the validity of these selective theories.

In *Drosophila*, SSD is regulated by the signaling pathways that control body size plasticity in response to developmental nutrition, in particular the IIS-pathway. This mechanism is consistent with the condition dependence hypothesis that explains SSD as a consequence of sex-specific selection for body size plasticity. If this hypothesis were correct, then we would expect (1) that the same loci underlie variation in SSP and SSD, and (2) that these loci lie in genes that are part of, or regulate, pathways that control growth in response to developmental nutrition, specifically the IIS pathway. However, our GWAS does not support these predictions: we found almost no overlap between candidate genes for SSD and SSP, and only few of them have previously been implicated in IIS regulation specifically, or nutritional signaling in general.

### GWAS Results Across Body Size, SSD, Plasticity, and SSP

We have previously found a weak but significant genetic correlation between sex-averaged body size and SSD, and a strong genetic correlation between SSD and SSP (14). Nevertheless, our GWAS revealed little overlap among candidate genes across these traits, and only two of the genes tested in our functional analysis demonstrably affected SSP. Both contained the SNPs with the two lowest *p*-values in our GWAS, and are regulators of gene expression, either through chromatin modification (*wda*) (31) or as a transcription factor (*Pdp1*) (32). Functional analysis of the remaining candidate genes revealed effects on sex-averaged plasticity, SSD, and sex-averaged body size, but not on SSP itself.

The lack of overlap among GWAS candidates, combined with the observation that functional manipulation of candidate SSP genes generally affected the other three phenotypes, may reflect the hierarchy among these traits. SSP is not an independent trait but instead emerges when SSD varies across environments. Similarly, SSD emerges when body size varies between sexes. Any genetic perturbation that alters SSP must necessarily also alter SSD, size plasticity, or body size in at least one sex and/or environmental condition.

Thus, genome-wide analyses may detect statistical associations with SSP through higher-order interactions (sex-by-diet-by-genotype), while functional disruption uncovers direct effects on growth regulation that manifest as differences in lower-order phenotypes such as SSD or plastic responses. In this way, the results of GWAS and functional analyses can appear contradictory but may actually be complementary, with GWAS highlighting small-effect QTLs specific to SSP and functional tests revealing a larger, more general, growth-regulatory role for the genes these QTLs inhabit.

### Candidate Genes and the Role of Canonical Growth Pathways

As noted above, several developmental studies have implicated the IIS pathway in regulating SSD and SSP in *Drosophila*. More generally, we might *a priori* expect genes that regulate variation in body size, SSD, size plasticity, and SSP to lie within pathways involved in growth regulation, sex determination or nutritional plasticity. Yet, with the exception of weak enrichment for growth regulators (**Table 2**), our GWAS found little evidence for the involvement of genes in the IIS or other canonical growth pathways.

One notable exception is *Pdp1,* one of our strongest GWAS hits, which is a basic leucine zipper transcription factor. The *Pdp-1* gene encodes multiple transcripts, one of which, *Pdp1ε*, is part of the transcription–translation feedback loops that maintain circadian rhythm in *Drosophila* (33–36). A second isoform, *Pdp-1γ*, is expressed in the fat body and appears to regulate lipid content (37). Deletion of *Pdp1* triggers non-cell-autonomous, nutrition-sensitive growth arrest that can be partially rescued by dietary sucrose (32). Further, *Pdp1* expression is upregulated under starvation conditions (32). These data support a role for *Pdp1* in coordinating cell proliferation and growth programs with nutrition, and, potentially, the regulation of sex-specific nutritional plasticity.

Nevertheless, the broader pattern in our study shows a general incongruence between QTLs for body size traits and known growth regulators, a theme that also emerges in other *Drosophila* GWAS on body and trait size. Vonesch et al. (30) conducted a GWAS on the size of the wings and the head, while Lafuente et al. (38) applied a GWAS to thorax and abdominal size and their thermal plasticity. Neither study found strong support for enrichment of growth-regulatory pathways, though both this study and Lafuente et al. found weak evidence for the involvement of Wnt- and Hippo-signaling in size plasticity. Surprisingly, only one candidate gene for variation in size was shared across all three studies: *jing*, a zinc-finger transcription factor involved in nervous system, tracheal, and imaginal disc development (39), but not directly implicated in growth regulation.

A similar pattern is not, however, seen in GWAS studies in other animals. In humans, GWAS for stature have revealed myriad variants, but enriched for genes in core growth pathways (e.g. Hedgehog/IHH, BMP/TGF-β) (40, 41) and growth-plate biology (42), and shows substantial overlap with Mendelian growth-disorder genes (40, 43, 44). In domesticated animals, GWAS has revealed causative variants in genes previously implicated in body size regulation, in particular the IIS pathway (45–49). The prevalence of growth regulatory genes in these GWAS may be explained by the high power of these studies and it is possible that more powerful GWAS on size traits in *Drosophila* will eventually uncover QTL in core growth regulatory pathways, albeit with smaller effect sizes. However, the loci identified so far do not support the hypothesis that natural variation in body size traits is controlled by genes known from developmental studies to regulate growth and plasticity.

### Private QTLs as Drivers of Trait Variation

Although the sizes of different body structures (e.g., head, wing, thorax, abdomen) are genetically correlated and co-regulated by shared developmental networks, both our GWAS and previous studies consistently identify loci with effects that are highly context-dependent and that impact only specific trait/environment /sex combinations within the trait hierarchy (e.g., body size, SSD, plasticity, SSP) (30, 38). This suggests that trait variation is driven by *private QTLs*: loci that affect variation in a general characteristic (e.g. growth) but only in a specific trait, sex, environment or some combination thereof (38)(38). Private QTLs represent a particularly stringent form of context-dependent QTLs (50), in which genetic effects are restricted to certain conditions rather than simply differing in magnitude. Evolutionarily, such private QTLs may facilitate adaptive change, as they may accumulate and maintain cryptic genetic variation that can be revealed in novel environments and allow selection to act on specific trait–sex–environment combinations without broad pleiotropic costs (51–53) although see (54).

### Mechanistic Models for Private QTLs

Our results suggest that private QTLs exhibit two layers of independence that limit their pleiotropy among hierarchical traits. First, they tend to occur in genes at the periphery of core signaling pathways. Second, within those genes, individual alleles have context-dependent effects. This structure is reflected in our data: few candidate QTLs fall within canonical growth pathways, and while the QTLs have narrow, context-specific effects, forced misexpression of the genes they inhabit reveals the gene’s more expansive pleiotropic potential.

Conceptually, there are two non-exclusive ways in which private QTLs may act in a context-dependent manner:

First, private QTLs may regulate pathways that integrate environment-, sex- and trait-specific information with processes underlying cell growth and proliferation. For instance, GxE effects that underlie variation in nutritional plasticity may arise through trait-specific changes in the sensitivity of the IIS pathway. This is observed in the reduced nutritional-sensitivity of male genital size in *Drosophila* (55) and the heightened sensitivity of male horn size in *Onthophagus* beetles (56). GxS (sex) effects that generate SSD may act by regulating tissue-specific responses to the sex-determination pathway, as seen in sex-specific horn size (57). GxT (trait) effects that cause trait-specific variation in size may impact patterning pathways, such as those producing wing–halter size differences in *Drosophila* (58).

Second, private QTLs may reflect condition-dependent activation of general processes such as metabolism, growth, or developmental timing. In this model, broad regulators (e.g. hormones, transcriptional co-factors, or chromatin remodelers) exert localized effects due to developmental context or epistasis (53), yielding QTLs that appear trait-, sex-, or environment-specific. For example, in *Drosophila* (and most animals), sex-biased chromatin accessibility means that some enhancers are accessible only in one sex (59). Consequently, genetic variants in these enhancers or in the transcription factors that bind to them, will have sex-specific effects, contributing to private QTLs. Similarly, transcription co-factors present only in one sex, tissue, or environmental condition may restrict the effect of genetic variation in the enhancers where they are recruited.

While the first mechanism reflects variation in the sensitivity of tissues or traits to shared upstream signals that communicate environmental-, sex-, or tissue-specific information, the second reflects variation in where and when regulatory elements are active, thus shaping the developmental context in which downstream pathways are engaged. Acting independently or in tandem, these mechanisms highlight how the phenotypic effects of genetic variation are gated by context, thereby generating private QTLs.

### The Mutational Target of Body Size and Future Directions

Further functional studies are necessary to distinguish among mechanisms for private QTL action. Our observation that functional manipulation of ostensibly private QTLs for SSP nevertheless impacts SSD, size plasticity, and body size suggests that at least some private QTLs lie in genes that regulate general biological processes rather than in exclusively trait-, sex-, or environment-specific growth. However, both mechanisms may act together, with some QTLs targeting canonical growth integration and others modulating general physiology.

What is clear is that body and trait size represent very large selective targets, regulated by a highly complex genetic network(60). As a consequence, the mutational target of body size is enormous, spanning not only growth pathways but also metabolism, developmental timing, tissue patterning, and physiological homeostasis. This breadth helps explain why GWAS on size traits in *Drosophila* consistently identify many loci of small effect (29, 30), often outside canonical growth regulators, and underscores why body size remains one of the most polygenic traits known (60–63). Because SSD and SSP arise from sex- and environment-specific modifications to these same developmental processes, their genetic architecture is similarly broad and distributed. The next challenge is to identify which of these loci actually contribute to evolutionary change in body size and its plasticity, SSD, and SSP, and to determine whether they act by modifying canonical pathways, altering their deployment in a context-specific manner, or by reshaping more general aspects of physiology and development.

## Methods

All data and the R code used to analyze them are provided on Dryad (DOI: 10.5061/dryad.8sf7m0d36)

### Fly Stocks

Flies come from the Drosophila Genetic Reference Panel (DGRP), consisting of more than 200 inbred *Drosophila melanogaster* lineages generated from a natural population in Raleigh, North Carolina, after 20 generations of full-sibmating (64). Each lineage discussed in this paper refers to a specific isogenic line in the DGRP with a unique genotype.

### Starvation treatment and Measurement

We collected data from 196 DGRP lineages, rearing flies according to established protocols (65, 66). Eggs were collected over a 12–20 hour period, placed in groups of 50 into 7 mL of standard cornmeal-molasses medium, and reared at 22°C until the starvation treatment was applied. To generate variation in body size, larvae were removed from food at precisely timed developmental stages and subjected to starvation either 0–24 h or 24–48 h before pupation. Since larvae naturally stop feeding ∼24 h before pupation (67), those removed 0–24 h before pupation were effectively fed ad libitum and are referred to as *fed* flies. In contrast, those removed 24–48 h before pupation were starved during the terminal growth period when nutrition affects adult size (68), and are referred to as *starved* flies. Starved larvae were transferred to empty vials with a wet cotton plug and kept there until pupariation. Pupae were then moved to individual 2.5 mL Eppendorf tubes (with a small puncture for gas exchange) to allow post-eclosion identification. After emergence, flies were sexed, and their associated pupal cases imaged and measured using semi-automated software from the Shingleton lab (69). Pupal case area (dorsal silhouette) was used as a proxy for adult body size (65, 66, 70). Flies were collected across nine temporal blocks, with five lineages repeated in all blocks as controls. We filtered out sex-fed/starved-lineage groups that had fewer than ten body size measurements, leaving fed-body sizes for 186 lineages and starved body sizes for 174 lineages.

### Lineage-Specific Phenotypes

All analyses were conducted in R (v. 4.0.3), and the script and data used in the analyses are provided on Dryad. Prior to analysis, all data were log transformed. We collected data from 15733 flies (7662 females and 8071 males; 8593 fed and 7140 starved) across 196 lineages.

We used a Mixed Linear Model (R package: *lme4*) (71) to generate Best Linear Unbiased Predictions (BLUPS) of fed and starved male and female in each lineage to calculate summary indices for subsequent GWAS. Specifically, for both the fed and starved flies we fit the model:

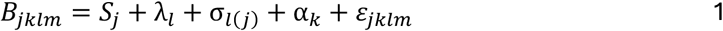

where *B* is body size, *S* is sex, and the model includes a random intercept *λ*_*l*_ for each lineage, a random sex-specific slope for sex *σ*_*l*(*j*)_, a random intercept for each block (α_*k*_), and residual error (*ε_j_*_*klm*_). We then extracted estimates of fed and starved male and female ^body size in each lineage^ 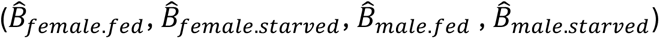.

Sex-averaged body size (fed or starved) was calculated as:

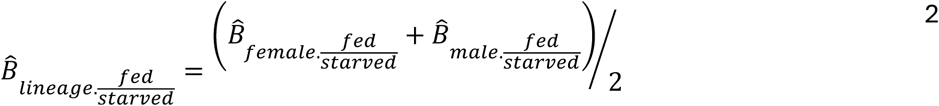

Sexual size dimorphism (fed or starved) was calculated as:

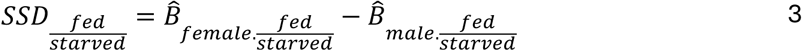

Nutritionally-induced size plasticity for each sex within a lineage was calculated as

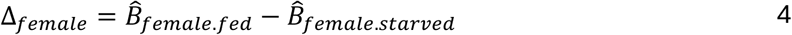

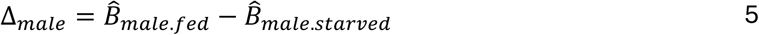

Sex-averaged plasticity was calculated as:

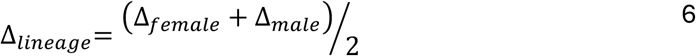

Sex-specific size plasticity for a lineage was calculated as

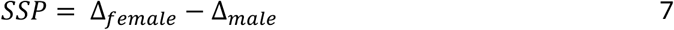

Because SSD (Eqn. 3) and SSP (Eqn. 6) are both calculated using *B̂*_*female.fed*_ and *B̂*_*male.fed*_ we *a priori* expect them to be correlated, which confounds subsequent GWA analyses. We therefore calculated an index of plasticity that was independent of *B̂*_*fed*_ by regressing *B̂*_*starved*_ on *B̂*_*fed*_ for males and females respectively, using OLS regression, and then used the residuals of the fit as an independent index of plasticity (Δ_*female.indep*_ and Δ_*male.indep*_). (See (14) for details). These indices were then used in Eqn. 7 to calculate independent sex-specific plasticity (*SSP*_*indep*_).

### Genome-Wide Association Study (GWAS)

We used the DGRP2 pipeline (https://quantgenet.msu.edu/dgrp/) to conduct a GWA analysis on: 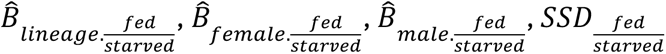, Δ_*lineage*_, Δ_*female*_, Δ_*male*_, *SSP*, and *SSP*_*indep*_. The pipeline tests for phenotypic difference between lineages containing major and minor alleles for 1,920,276 genetic markers (SNPs, short deletions and insertions) across the *Drosophila* genome. The pipeline corrects for the presence/absence of *Wolbachia* and five major inversions present within the DGRP lineages [In(2L)t, In(2R)NS, In(3R)K, In(3R)P and In(3R)Mo]. The analysis also corrects for genetic relatedness by incorporating covariates that correspond to the major principle components of the relationship matrix.

The DGRP2 GWAS pipeline is computationally efficient because it uses lineage-level summary values to conduct the GWA analysis on a phenotype; that is, for each genetic marker it fits a linear model where *n*≈196. These values (e.g. *B̂*_*female.fed*_, *B̂*_*female.starved*_, *B̂*_*male.fed*_, *B̂*_*male.starved*_), however, do not capture the uncertainty of measurements within each lineage, and, in our study at least, are based on variable number of individuals (between 10 and 290). This can generate anticonservative estimates of statistical significance for association tests using indices based on these values (72, 73), increasing the likelihood of Type I error.

To reduce the possibility of false positives, we conduct a subsequent mixed linear model (MLM) GWA analysis using all the collected data, which in our study comprised >26,000 body size measurements. Specifically, we fit the model:

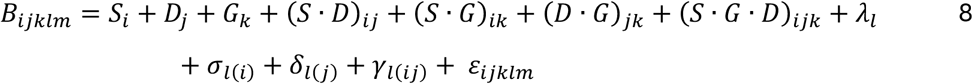

where *G* is the bivariate allelic marker, *D* is diet treatment (fed or starved) and each lineage (*l*) has a random intercept *λ*_*l*_, a random slope for sex *σ*_*l*(*i*)_, a random slope for diet *δ*_*l*(*j*)_ and a random slope for sex-by-diet *γ*_*l*(*ij*)_ (subscripts indicate levels within factors). Here, (*S* · *G* · *D*)_*ijk*_ captures the effect of genotype on SSP. By definition, a significant (*S* · *G* · *D*)_*ijk*_ interaction meant that the genotype affects plasticity (at least in one sex), SSD (at least in fed or starved flies) and body size (at least in one sex in fed or starved flies).

While powerful, this approach is computationally slow (the analysis of each SNP takes ∼1s on a powerful laptop computer). Consequently, we only fit the MLM GWA analysis to markers likely to be associated with variation in our phenotype, restricting it to the 10,000 SNPs with the lowest *P-*values from the DGRP2 GWAS of *SSP* and *SSP*_*indep*_, which, due to common hits between the two DGRP2 GWAS, comprised 12,933 SNPs. As a control, we repeated the analysis on 12,933 SNPs randomly selected from the remaining pool.

Prior to the MLM GWAS we removed block effects by regressing body size on block and using the residuals in the analysis. As for the DGRP2 pipeline, we corrected for the presence/absence of *Wolbachia*, inversion status and population structure, by substituting genotype (*G*) in Eqn 8. with *Wolbachia* status (*W*), inversion status (*I*) [the five inversions above, plus In(2R)Y1, In(2R)Y5, In(3L)P, In(3L)M, In(3L)Y, In(3R)C], and the first 20 principal components (*PC1–20*) of the genetic distance matrix (available at http://dgrp2.gnets.ncsu.edu/data.html). These effects were tested first as interaction effects, then as additive effects, and removed if found to be non-significant.

The same two-step method was used to conduct GWAS on 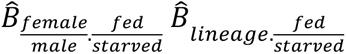, 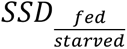, Δ_*male*_, Δ_*female*_, and Δ_*lineage*_, but by fitting different MLMs (**Table 3**). Prior to analysis we again removed block effects, and included *Wolbachia*, inversion status and the principle components of the genetic distance matrix as covariates in the model, if they had a significant effect on body size.

For each MLM GWAS we identified candidate SNPs using the commonly-used *p*-value of < 1×10⁻⁵ (29, 38, 64, 74–76), and annotated the candidates using FlyBase release 5.49.

### Versatile Gene-based Association Study (VEGAS)

We used a VEGAS-style approach to test whether multiple SNPs within the same gene carried an aggregate signal of association, while accounting for gene size and linkage disequilibrium (LD) structure (76). For each gene, we converted the MLM GWAS *p*-values of its constituent SNPs into equivalent 1-degree-of-freedom chi-square statistics, by taking the inverse chi-square quantile, 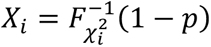, where 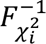 is the inverse cumulative distribution function of chi-square. Here, smaller *p-*values correspond to larger chi-square values. We then summed these values across all SNPs in the gene to obtain an observed gene-level statistic, ignoring SNPs that had no variation across lineages. To account for gene length and the non-independence of SNPs due to LD, we estimated the correlation structure among SNPs within the gene from the genotype matrix. This correlation matrix was adjusted to be numerically stable and positive-definite. We then generated a null distribution for each gene by sampling 10,000 replicates of multivariate normal vectors with the same LD structure, squaring their values to mimic chi-square statistics, and summing them across SNPs. The empirical gene-level *p*-value was defined as the proportion of simulated statistics that were smaller than or equal to the observed statistic. Candidate genes were selected based on the threshold *p*-value of < 1×10⁻⁵.

### Enrichment Analysis

We conducted enrichment analyses on candidate genes in three ways:

First, we conducted a GO and KEGG analyses of candidate genes from the MLM GWA and subsequent VEAGS analysis using the *clusterProfiler* package in *R* (77).

Second, we performed a permutation-based test to evaluate whether genes in growth-regulatory or nutrient-signaling pathways were enriched for SNPs with low *p*-values in our MLM GWAS. For each of four phenotypes (*B̂*_*lineage.fed*_, *SSD*_*fed*_, Δ_*lineage*_, and SSP), we began with the ∼10,000 SNPs that had the smallest *p*-values in the initial DGRP2 GWAS and that were subsequently tested in the MLM GWAS. These SNPs were mapped to a list of 331 canonical growth genes and, separately, to 70 nutrient-signaling genes (**Supplementary Table S4**). Each gene was summarized by the minimum MLM p-value across its SNPs. As a test statistic, we used the mean of these per-gene minima within each gene set. To generate a null distribution of the test statistic, we randomly sampled the same number of genes from the ∼10,000 SNPs and calculated the mean per-gene minimum p-value for each sample, with 10,000 replicates. Enrichment was assessed as the empirical percentile (and corresponding one-sided empirical *p*-value) of the observed mean relative to the null distribution.

Third, we conducted a VEGAS analysis of the same growth-regulatory and nutrient-signaling genes, as described above.

### Functional Validation

Based on the results of our MLM GWA analysis, we selected a list of candidate genes to test for their effect on SSP. We prioritized markers within the coding or regulatory regions of annotated genes with known functions. To validate gene function, we manipulated the expression of twenty candidate genes ubiquitously using act-GAL4 to drive UAS-RNAi constructs from either the Drosophila Transgenic RNAi Project (TRiP) collection or the Vienna Drosophila Resource Center (VDRC) collection. For each gene we crossed flies homozygous for the UAS-RNAi construct with *yw;act-GAL4/CyO*, which was generated by backcrossing *act-GAL4* into a *w^1118^* isogenic stock for seven generations and then crossing with the co-isogenic *w^1118^;+/CyO*. In each cross, control flies carried *CyO* while experimental flies did not. Fed and starved progeny from each cross were generated as described above, and their pupal cases sizes measured as a proxy for body size. To test the effects of gene knockdown, we fit the following model:

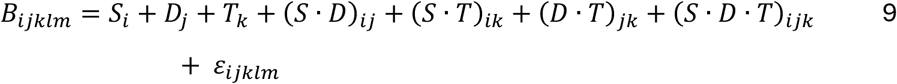

where *T* is fly type (experimental or control). Here, *S* · *G* · *T* captures the effect of knockdown on SSP, *D* · *T* captures the effect on plasticity, and *S* · *T* captures the effect on SSD. Where necessary, the model was refit after removing the least significant higher-order interaction until only the simplest model remained.

