## Supplementary Figure for "Private QTLs and the Genetic Architecture of Hierarchical Size Traits: From Body Size to Sex-Specific Plasticity"

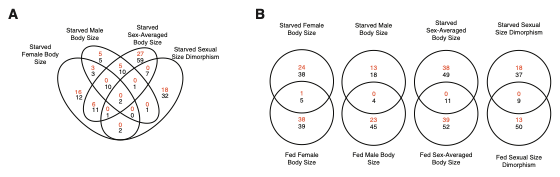

**Supplementary Figure S1.** Overlap of candidate SNPs (red) and genes (black) for different size-related phenotypes. (A) Overlap between male, female and sex-averaged body size and SSD in starved flies. (B) Overlap in male, female, and sex-averaged body size and SSD between fed and starved flies. Candidate SNPs were identified by MLM GWAS and candidate genes were identified by VEGAS.

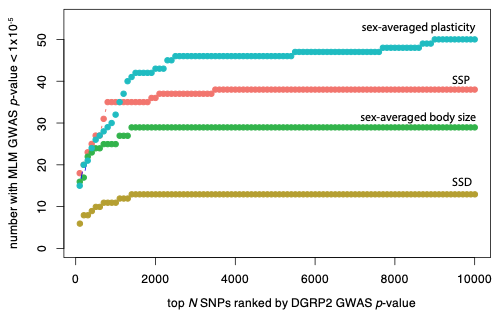

**Supplementary Figure S2.** The accumulation of MLM GWAS hits with the inclusion of increasingly large number of SNPs, ranked by their DGRP2 GWAS *p-*value. For most traits, the accumulation plateaus after ~4,000 SNPs, suggesting that conducting an MLM GWAS on the 10,000 SNPs with the lowest DGPR2 GWAS *p-*value provides a sufficiently inclusive set to capture all MLM GWAS hits (*p*<1x10^-5^).
